# Bridging Genomic Gaps: The Pan-Grass Syntenic Gene Set in Grass Crop Evolution

**DOI:** 10.1101/2024.11.12.623179

**Authors:** Henrique M Dias, Guilherme YA Sagawa, Vladimir J Torres-Rodriguez, Ravi V Mural, James C Schnable, Marie-Anne Van Sluys

## Abstract

Understanding orthologous and homeologous relationships among genes can inform our understanding of how evolution and artificial selection have shaped the genomes of related species to adapt to different niches and produce different phenotypes. High-confidence information on orthologous relationships can also increase the impact of the functional characterization in one species by ensuring this information is available to researchers studying orthologs in other species. The grasses are ecologically and economically critical plants, and many genes are conserved in syntenic blocks between species. Here we leverage that conserved gene order to create and describe the Pan-Grass Syntenic Gene Set (PGSGS), a curated database of orthologous and homeologous relationships among the genes of 17 grass species, including both major and orphan crops as well as wild grasses. As a demonstration of the utility of the PGSGS, the dynamics of chromosome structure and gene content changes after polypoidy in several grass lineages are analyzed. This dataset serves as a seminal foundation for improving our understanding of genome evolution subsequent to polyploidy, as well as encouraging and enabling integration of functional genetic and functional genomic information across the grasses, the original promise of the “grasses as a single genetic system” model.

**SIGNIFICANCE STATEMENT:** The Pan-Grass Syntenic Gene Set (PGSGS) leverages our understanding of the genomic background of major grass species, including rice, maize, sorghum, wheat, barley, millet, and sugarcane. It represents a milestone for deciphering the genetic foundations of essential traits such as yield, disease resistance, and environmental adaptability - key factors contributing to agriculture and food security. Scrutinizing pan-syntenic genes unveils the intricate interplay of diverse gene acquisition and loss mechanisms, potentially speeding molecular breeding knowledge transference. The PGSGS emerges as a valuable resource for gene annotation, with implications for improving grass crops.

## INTRODUCTION

Approximately 50% to 60% of the world’s calorie intake comes from grain crops (grasses) including the major cereal crops, such as rice, wheat, and maize, as well as minor and orphan crops such as pearl, foxtail, and proso millet (1). Beyond grains, other grass species also play key roles in agriculture as sources of sugar (sugarcane), forages for livestock, and feedstocks for bioenergy production (2). Wild grass species are ecologically dominant in vast areas of most continents, and a broad set of molecular, anatomical, and physiological adaptations has allowed the members of this clade to thrive in disparate environments (3). Although grasses exhibit a broadly conserved and collinear genomic architecture, they harbor extensive genetic diversity, encompassing variations in nucleotides (single-nucleotide polymorphisms), differences in gene structure, and the presence or absence of specific genes responsible for particular traits among different species (2). Grasses exhibit a wide range of photosynthetic and water use efficiencies (4-6); different grass species utilize C3 photosynthesis or three different types of C4 photosynthesis, and different subclades within the grasses exhibit different capacities to adapt to and thrive in stressful and resource-poor environments (7).

Despite their diverse morphologies and ecological niches, the genomes of most grasses exhibit a high degree of genetic and genomic collinearity (8-10). In a typical grass genome, approximately half of all annotated gene models will be conserved at syntenic locations related to other grasses, but these syntenically conserved genes will include the vast majority—greater than 90%—of genes linked to phenotypic variation (11). Both rice and sorghum are diploid relative to the most recent common ancestor of the grasses, and the reference genomes of these two species are widely used in grass comparative genomic analyses. However, many other economically important grass species are either mesopolyploids (maize) or neopolyploids (wheat, sugarcane, switchgrass, miscanthus). These whole genome duplications can create two or more syntenic intervals that are each co-orthologous to a single common region in the genome of an unduplicated grass such as sorghum or rice (12; 13). Distinct patterns of expression and/or function can be observed across different subgenomes in polyploid grasses (14; 15). From this standpoint, comparative genomic analysis is particularly fruitful in the grasses, as synteny can be extremely informative both on a single gene level regarding functional data from orthologs in other species and on a genome-wide level in understanding the dynamics of genomic evolution post-polyploidy (16). Various publicly accessible tools, including Phytozome (17), CoGe (18; 19), PLAZA (20), Gramene (21), Ensembl Plants (22), and others, can be employed for automated comparative genomics analyses between or among grass genomes and those of other plant species.

However, in many cases, these tools do not capture the subgenome identity of individual gene copies in polyploid species. Here we employ a previously published methodology to update a widely cited and reused syntenic gene set from seven years ago (15), incorporating syntenic analysis of additional agriculturally, ecologically, or evolutionarily relevant grass species. The result is the Pan-Grass Syntenic Gene Set (PGSGS). The PGSGS provides an optimized approach for exploring the evolution of individual genes across diverse grass species groups. This database encompasses 17 genomes that have undergone one or more WGD, including both diploid and polyploid genomes. As a demonstration of the potential applications of this dataset, we conduct a case study investigating the evolution of core grass genes.

## MATERIAL AND METHODS

### Identification of Syntenic Orthologous Genes

A set of seventeen grass genomes representing fifteen species, all available as part of Phytozome v13 (17), were employed for analysis in this paper (Table S1). For each genome, the “cds_primaryTranscriptOnly” fasta file and GFF annotation files were downloaded from Phytozome for downstream analyses. The workflow for pairwise identification of syntenic orthologous genes between the sorghum reference genome and the other sixteen genomes is summarized in Figure S1. LASTZ pairwise comparisons were performed using the primary transcript CDS sequences from the species of interest as query and the primary transcript CDS sequences of *Sorghum bicolor* BTx623 v5.1 as reference. LASTZ was run with a seed of match12, a minimum identity of 70%, and a minimum coverage of 50%. The output of LASTZ was employed to identify syntenic blocks between sorghum and the genome of interest using QuotaAlign (23) with pairwise comparison-specific sets of parameters given in Table S1. Finally, we polished the raw QuotaAlign gene pairs, selecting the gene showing the greatest sequence similarity (as determined by LASTZ alignment score) within the window from 20 genes downstream/upstream of the predicted location (15).

### Additional Bioinformatics Analyses

To elucidate the functional significance of the identified syntelog gene set, functional enrichment analysis was performed. Gene Ontology (GO) enrichment analysis was conducted using the package in R following a workflow by Bonnot et al. (24). The analysis focused on categorizing genes based on their biological process. Significantly enriched GO terms were identified using a Fisher’s exact test, P < 0.05 after False Discovery Rate (FDR) correction. The PGSGS was enriched using EggNOG v5.0 (25).

## OVERVIEW OF THE PGSGS

### A Robust Synteny-Based Identification of Evolutionary Relationships Among Grass Genes

The use of synteny information to identify orthologous and homeologous relationships among genes is robust to changes in evolutionary rate, pseudogenization, and other factors which can disrupt or cause misleading results in purely tree-based methods. A number of tools exist for identifying syntenic regions within or across genomes given the set of known homologous relationships among genes (18; 23; 26). However, in many cases, these tools can conflate orthologous and homeologous syntenic regions. We adopted a three-stage pipeline, employing LASTZ for the identification of homologous genes (27), QuotaAlign for identification of syntenic regions and the specific identification of orthologous syntenic regions (23), and a final polishing step to improve orthologous gene assignment, particularly for genes with multiple local duplicate copies within the syntenic regions originally identified by QuotaAlign (15). This is particularly crucial in regions where there are tandemly duplicated arrays of genes or proximal paralogs that have undergone independent evolutionary trajectories over millions of years.

The effect of the polishing process is illustrated in Figure 1. In order to speed genome-wide comparisons, QuotaAlign collapses sets of homologous genes located within the same genomic intervals and considers only one gene per group when defining genome-wide synteny relationships (Figure 1A). In some cases, these collapsed genes are recent tandem duplications, but in other cases, they may represent genes which have subfunctionalized and evolved independently for long time periods. In the polishing step, the global synteny information from QuotaAlign is used to identify the predicted region syntenic orthologs should be present in, while the raw alignment scores for all possible pairwise gene comparisons are used to identify pairs of the most similar genes within these syntenic intervals (Figure 1A). The result is an increase in the number and accuracy of syntenic orthologous gene assignments (Figure 1B). In our case, using sorghum as a reference, this polishing step typically increased the number and accuracy of syntenic gene assignments by approximately 10% for species within the panicoid grass subfamily and by a smaller amount in most distantly related grasses outside the panicoid subfamily (Figure 1B).

**Figure 1.**
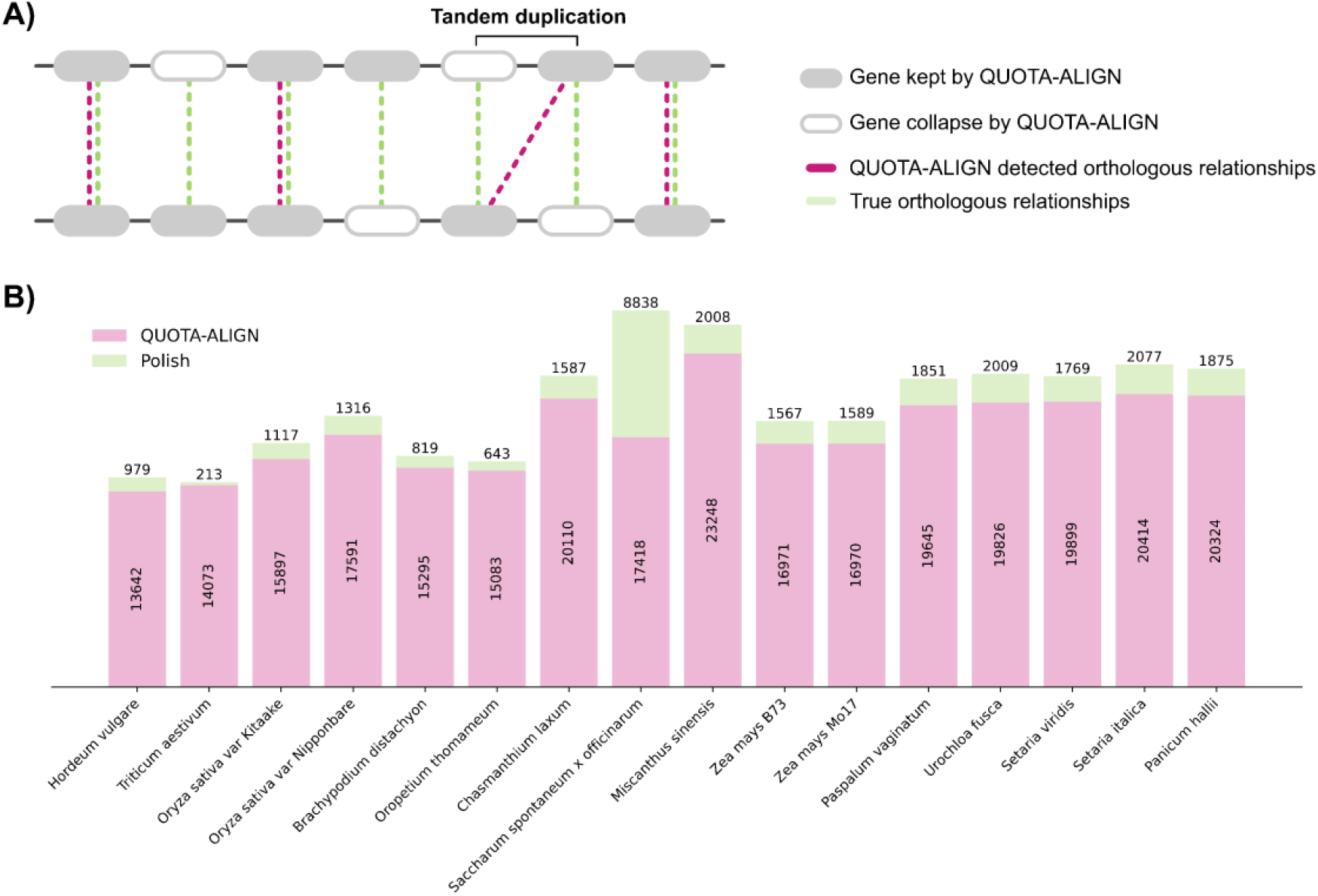
Schematic representation of a sequence homology-based approach to identify syntenic duplicated genes. Pink dashed lines depict the identification of orthologs through pairwise comparison between Genome A and B using the QUOTA-ALIGN method (15). Green dashed lines illustrate the refinement of paralog identification arising from duplications within the same genomic environment, employing a sequence homology-based approach.

### Development of a Database of Grass Orthologous Relationships

We selected a total of sixteen additional grass genomes, representing fourteen additional grass species, for syntenic comparison to our sorghum anchor sequence. We preferentially selected genomes with high N50/L50 metrics for genome quality, accurate gene annotations, and high BUSCO scores (Supplementary Table S1); however, several genomes with complex histories of duplication and/or more fragmented assemblies were also included based on evolutionary position and/or economic significance. This dataset represents both a significant expansion in the number of species captured from earlier pan-grass syntenic datasets (28) but also employs more recent gene assemblies and gene model IDs for many of the genomes included in the previous synteny dataset (Figure 2).

**Figure 2.**
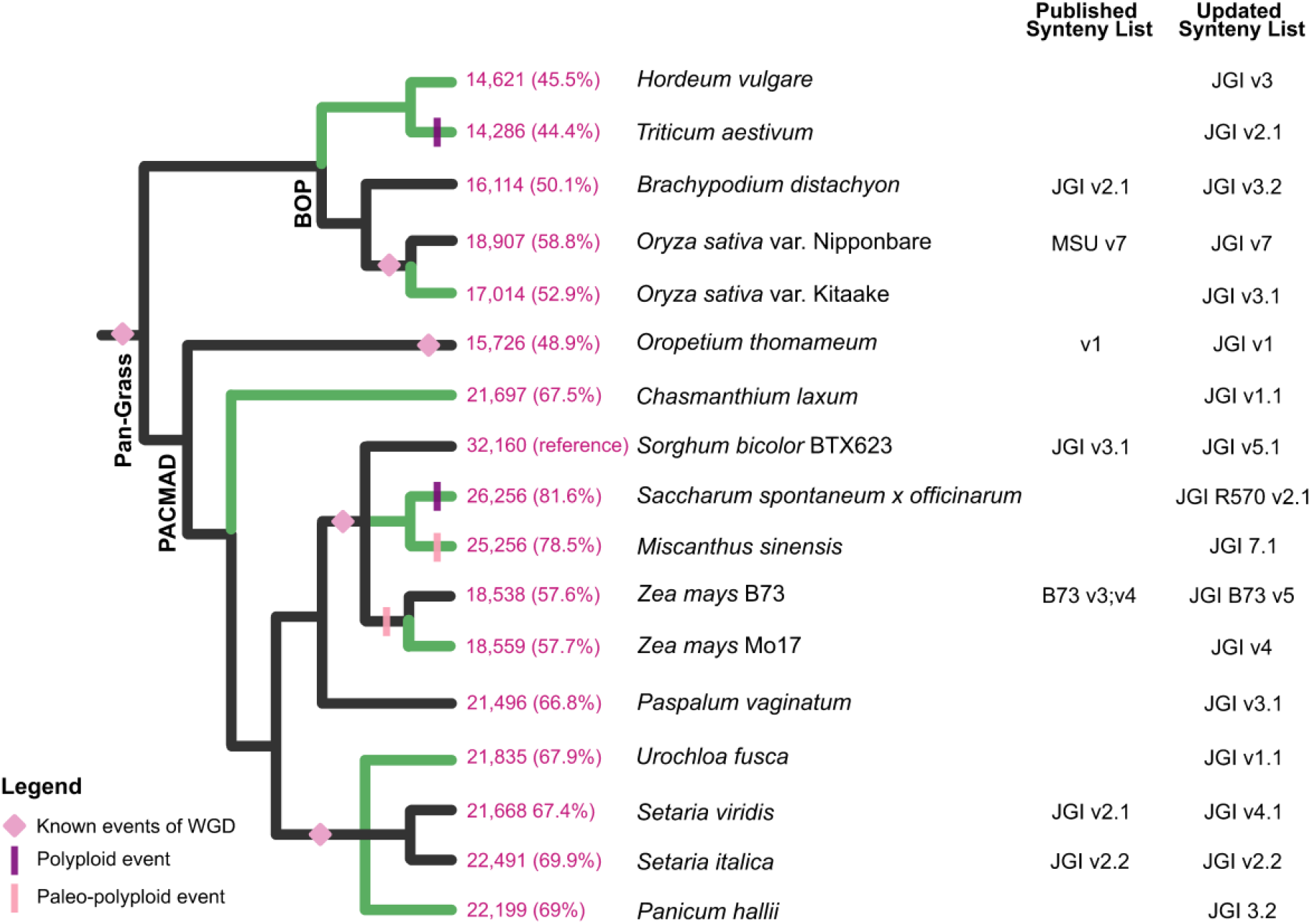
Phylogenetic relationships among genome releases of sequenced grass species and syntelog gene counts within the Pan-Grass Syntenic Gene Set (PGSGS). The PGSGS, with a Sorghum-based foundation, is presented, showcasing syntelog counts in pairwise comparisons. Absolute and relative syntelog numbers are depicted in pink, while green denotes genome versions added in the PGSGS update. The earliest genome versions were updated from the initial PGSGS version (https://doi.org/10.6084/m9.figshare.7926674.v1). Pink diamonds symbolize whole-genome duplication (WGD) events referenced by Zhang et al. (40), and polyploidy and paleo-polyploidy events are depicted by dashes.

Out of 32,160 gene models annotated in v5 of the *Sorghum bicolor* genome, 27,567 were linked to a syntenic ortholog in at least one other grass species. Each set of grass genes consisting of one sorghum gene and its syntenic orthologs in other grass species was assigned a unique PGSGS_ID (Supplementary Table S2). Roughly twenty-five thousand sorghum genes could be linked to syntenic orthologs in other core Andropogoneae species; eighteen to twenty-two thousand sorghum genes could be linked to syntenic orthologs in panicoid grasses outside the core Andropogoneae; and roughly fourteen to eighteen thousand sorghum genes could be linked to syntenic orthologs in grass species outside the panicoid subfamily (Figure 2). The reasonably slow decay in the number of conserved syntenic orthologs when comparing more distantly related grass species suggests a single anchor genome may work well for most cases in the grasses. However, it seems likely an independent analysis conducted using rice as a reference genome might identify several thousand additional syntenic orthologs specific to the BOP clade of grasses (29).

### Unraveling the Dynamics of PGSGS

As data has been gathered from greater numbers of individuals within many plant species, it has become apparent that many plants possess both a core gene set which is shared among the genomes of all individuals within the species and a set of non-core genes carried in the genomes of some, but not all, individuals within the species (30). A similar approach can be taken to defining core and non-core gene sets among clades of related species. We computed the non- and core-gene sets across the 17 grass genomes used in our study. The core set was defined as genes found in at least 15 genomes. The size of the combined core and non-core gene sets among these grass species continued to expand with the inclusion of additional genomes, indicating its open nature (Figure 3A). The size of the PGSGS-core rapidly declined as more genomes were added, particularly when considering the phylogenetic distance relative to sorghum (Figure 3B). In addition to the overall core gene set, we computed a second gene set specifically for the PACMAD group of species, which are more closely related to sorghum (Figure 2). The PACMAD-core consists of 2,266 syntelogs unique to this group, and shares 9,689 syntelogs with the PGSGS-core, making a total of 11,956 syntelogs. Conversely, the PGSGS-core includes 1,934 syntelogs (out of a total of 11,624 syntelogs) that are not part of the PACMAD-core.

**Figure 3.**
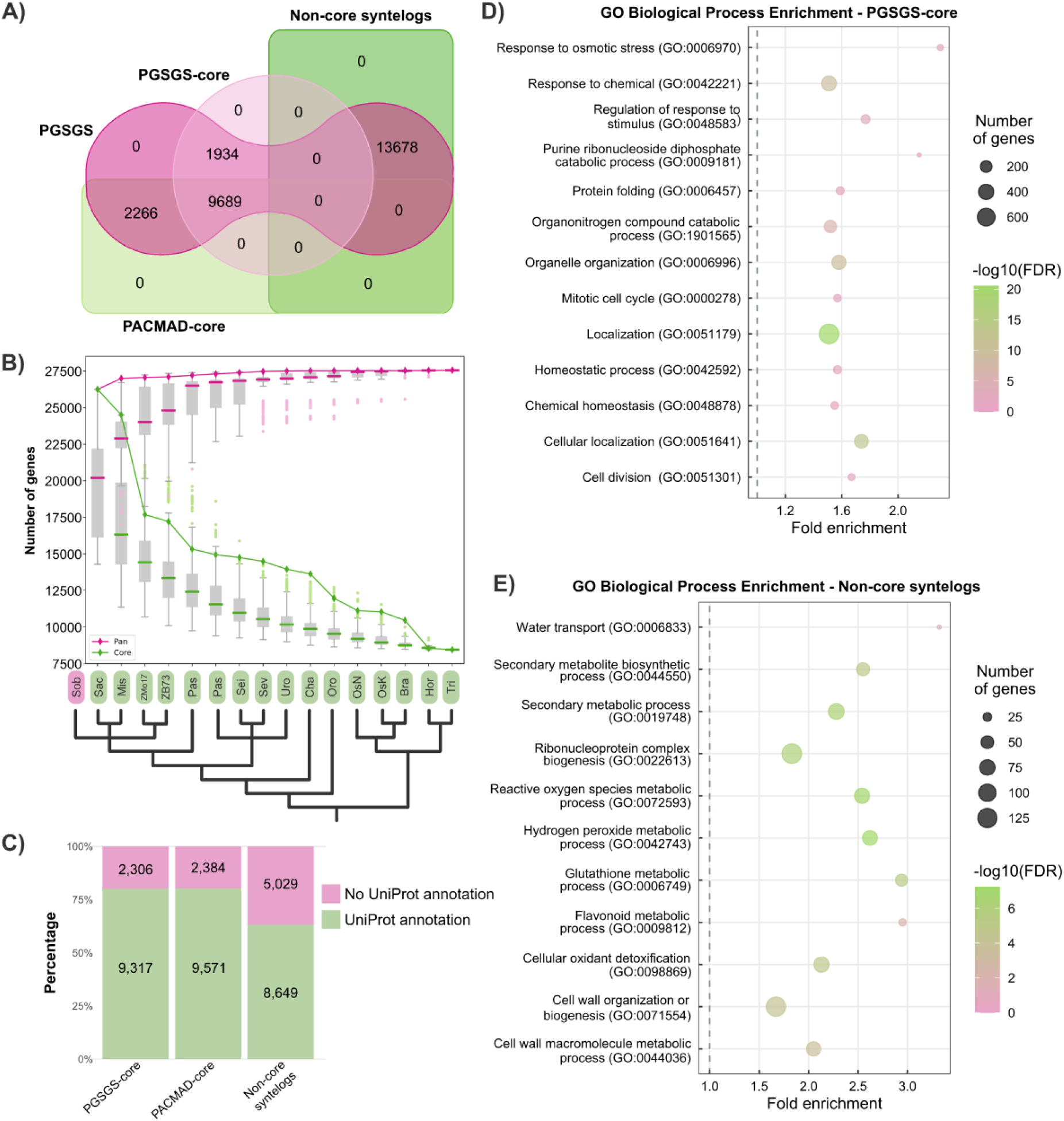
Pan-Grass Syntenic Gene Set (PGSGS) overview. **A)** The Venn diagram delineates the PGSGS, PGSGS-core, PACMAD-core, and non-core syntelogs. **B)** The PGSGS size increases for each included genome, with grey hues representing the 25th and 75th percentiles, while pink and green lines depict pan- and core-syntelogs median, respectively. Pink and green diamonds correspond to the exact number of syntelogs added by species following the phylogenetic tree. **C)** The number (indicated in the bars) and fraction of PGSGS-core, PACMAD-core, and non-core syntelogs containing annotated UniProt entries. **D-E)** Functional enrichment analysis of PGSGS-core and variable syntelogs, with significance (FDR < 0.05) and effect size (log2 fold change ≥ 1.5).

The pan-grass core gene set includes a notably higher proportion of gene models with annotated protein domains than does the non-core syntelog set. 80.2% of core syntelogs and 63.2% of non-core syntelogs are associated with a Uniprot annotation (Figure 3C). The pan-grass core set was enriched for genes annotated as involved in cell division (GO:0051301), cellular localization (GO:0051641), chemical homeostasis (GO:0048878), homeostatic process (GO:0042592), localization (GO:0051179), mitotic cell cycle (GO:0000278), organelle organization (GO:0006996), organonitrogen compound catabolic process (GO:1901565), protein folding (GO:0006457), purine ribonucleoside diphosphate catabolic process (GO:0009181), regulation of response to stimulus (GO:0048583), response to chemical (GO:0042221), and response to osmotic stress (GO:0006970) (Figure 3D), while the non-core syntelog set was enriched for cell wall macromolecule metabolic process (GO:0044036), cell wall organization or biogenesis (GO:0071554), cellular oxidant detoxification (GO:0098869), flavonoid metabolic process (GO:0009812), glutathione metabolic process (GO:0006749), hydrogen peroxide metabolic process (GO:0042743), reactive oxygen species metabolic process (GO:0072593), ribonucleoprotein complex biogenesis (GO:0022613), secondary metabolic process (GO:0019748), secondary metabolite biosynthetic process (GO:0044550), and water transport (GO:0006833) (Figure 3E and Supplementary Table S3).

### Defining Subgenomes and Homologous Chromosomes

Polyploidy, or whole genome duplication, is common in plants (31) and dramatically reshapes genome organization and gene content and may have significant impacts on evolutionary processes and adaptive characteristics (32; 33). Understanding the genetic mechanisms governing polyploidy in plants remains a significant challenge (34). Completing the PGSGS required a comparison of the sorghum genome to multiple mesopolyploid and neopolyploid species including *Zea mays, Miscanthus sinensis, Saccharum officinarum x spontaneum*, and *Triticum aestivum*. In recent polyploids such as *Triticum aestivum* (wheat), the subgenome identity of each chromosome is known a priori, while in less evolutionarily recent polyploidy/WGDs such as *Zea mays* (maize), subgenome identity can potentially be inferred from differential evolution of homeologous genomic segments (35). The most complex case of the polyploid genomes examined here was *Saccharum officinarum* x *spontaneum* (sugarcane). Sugarcane is a complex polyploid crop characterized by aneuploidy, resulting in irregular chromosome counts, typically displaying six to eight sets of chromosomes (2n = 6-8x), resulting from elaborate hybridization among diverse species within the Saccharum genus (36; 37). The complex, asymmetric chromosome composition resulting from polyploid *Saccharum* hybridization complicates the inference of syntelogs across homologous chromosome sets (see Figure 4, showing the aneuploidy phenomena between homologous chromosomes). These results are further supported by the recent publication of the R570 sugarcane genome (38). This scenario adds a layer of complexity to syntenic ortholog inference. Nevertheless, we have identified a robust set of homeologous genes across the homologous chromosomes of sugarcane (*Saccharum officinarum* x *spontaneum* var. R570) (39), including chromosomes A-G and the additional recombined chromosome OS (Supplementary Table 2).

**Figure 4.**
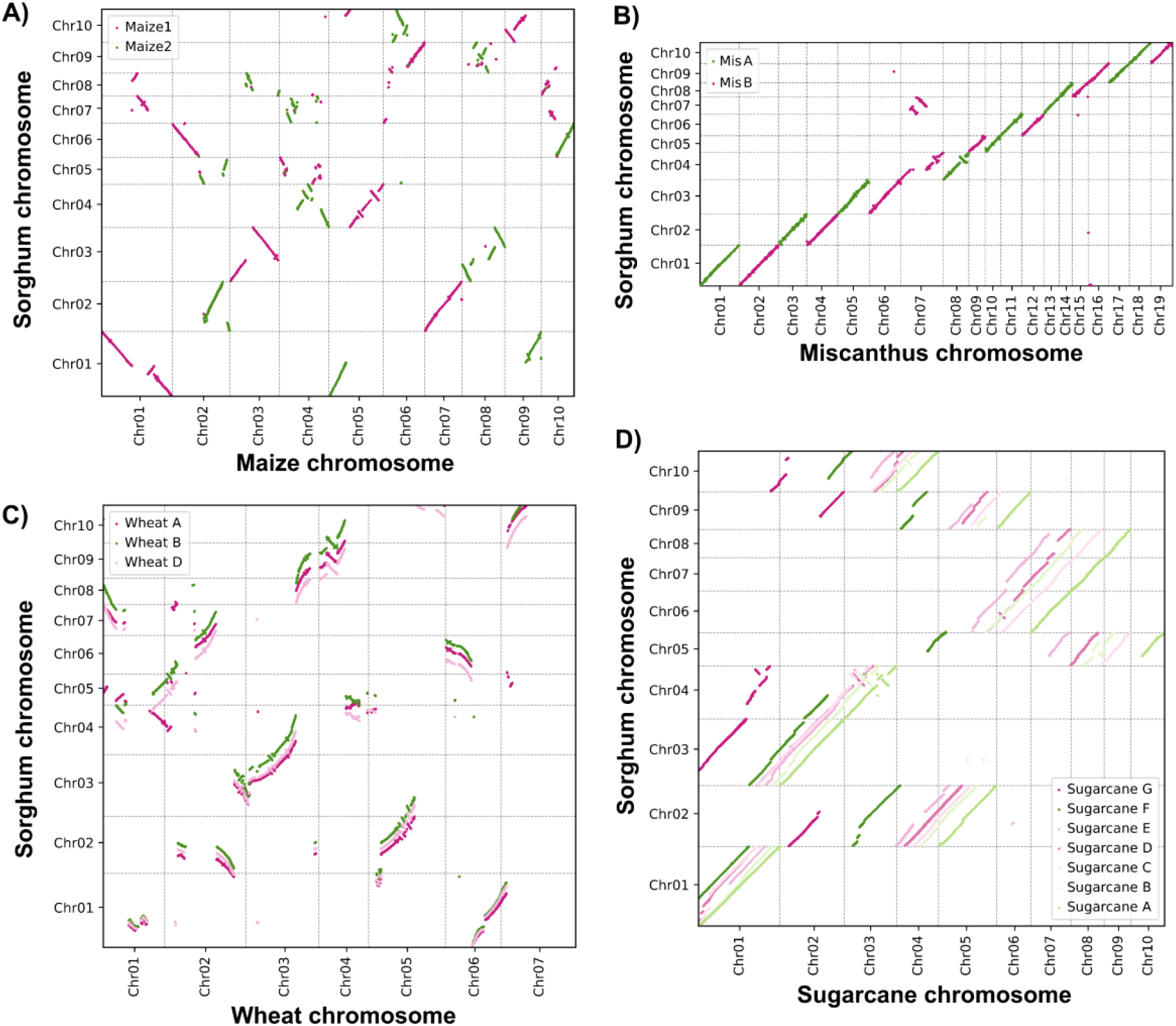
Dot plot illustrating syntelog clusters between *Sorghum bicolor* and paleo- and polyploid grass genomes. **A)** Genome collinearity analysis between *S. bicolor* and *Z. mays*, indicating maize 1 and maize 2 subgenomes. **B)** Genome collinearity analysis between *S. bicolor* and *M. sinensis*, highlighting MisA and MisB subgenomes. C) Genome collinearity analysis between *S. bicolor* and *T. aestivum*, illustrating wheat homologous chromosomes A, B, and D. **D)** Genome collinearity analysis between *S. bicolor* and *S. spontaneum* x *officinarum* var. R570, showcasing sugarcane homologous chromosomes A, B, C, D, E, F, and G. Different-colored dots represent subgenomes or homologous chromosomes.

### PGSGS Analysis Reveals Conserved Syntelogs and Species-Specific Adaptations

As a demonstration of how the PGSGS can aid research into genomic evolution, we analyzed a comprehensive matrix containing the total set of syntelogs (27,567) across various grass species. This analysis focused on genomes and homologous chromosomes, specifically examining the presence or absence of synteny for individual genes (Figure 5). The genomic organization of syntelogs across different grass species revealed intriguing patterns of similarity and divergence, particularly among subgenomes and homologous chromosomes. We identified a subset of highly conserved genes across multiple species, forming clusters of core syntelogs. Conversely, several genes exhibited species-specific synteny patterns. For instance, unique synteny profiles were observed in Sorghum bicolor and other grasses, suggesting species-specific genomic adaptations and possible functional specialization of non-core syntelogs.

**Figure 5.**
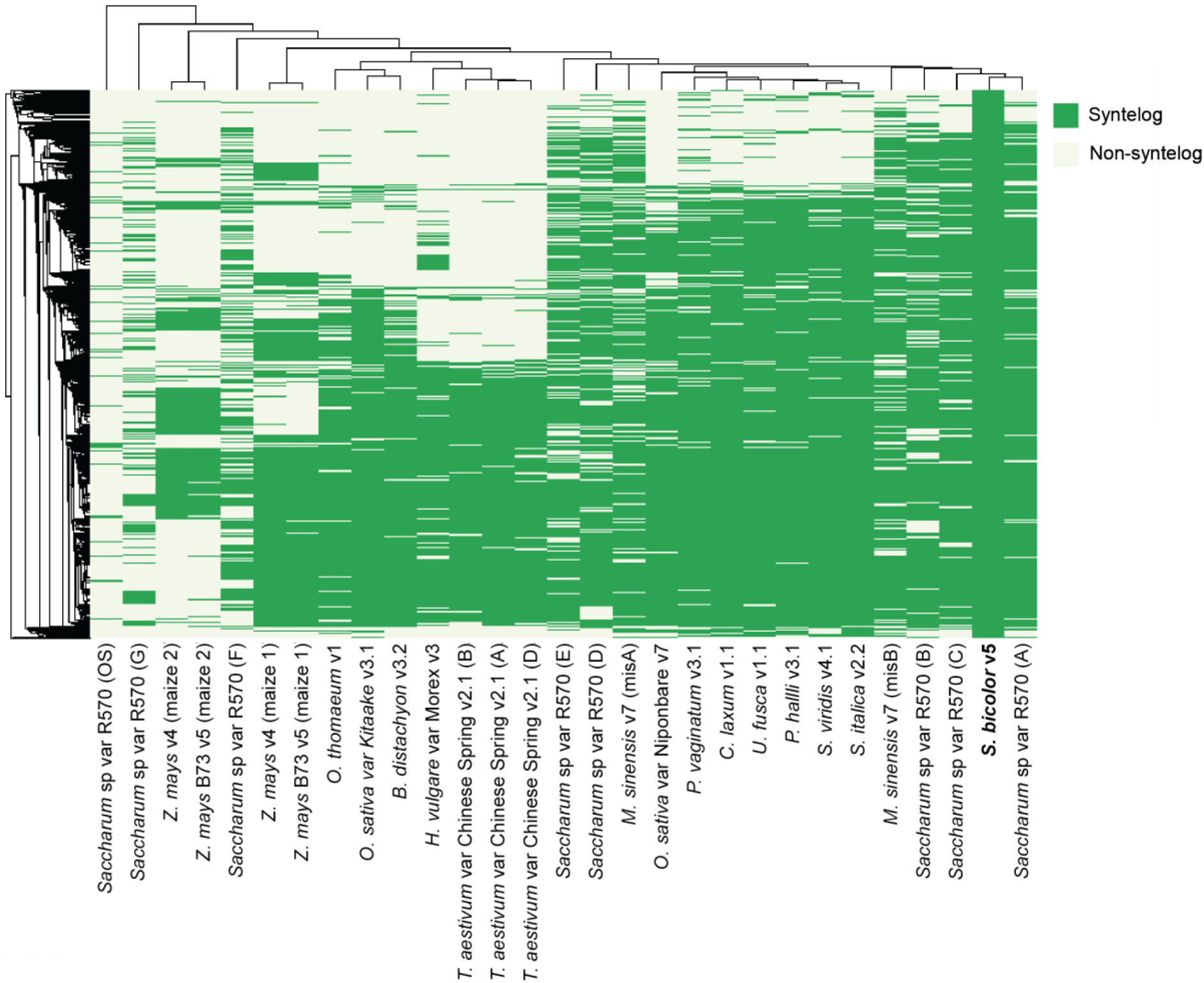
Synteny Presence-Absence Heatmap Highlighting Genomic Variation Across 17 Grass Species. This heatmap illustrates the synteny presence-absence across multiple genomes and chromosome homologs for 17 grass species. Each column represents a different genome or chromosome homolog, and each row corresponds to a gene. The color scale ranges from 0 (absence, white) to 1 (presence, dark green). Dendrograms indicate hierarchical clustering of genomes and genes, revealing conserved and divergent synteny patterns. Hierarchical clustering to identify patterns and group similar samples in the binary matrix dataset was performed using the Jaccard method with the pheatmap package in R (41). ‘Syntelog’ indicates presence, and ‘Non-syntelog’ indicates absence of synteny. The genomes represented are listed in Supplementary table 2 and are as follows: S.bicolor v5, H.vulgare Morex v3, T.aestivum_cv._ChineseSpring v2.1 (A); T.aestivum_cv._ChineseSpring v2.1 (B); T.aestivum_cv._ChineseSpring v2.1 (D), O.sativa Kitaake v3.1, O. sativa Nipponbare v7.0, Brachypodium distachyon v3.2, O.thomaeum v1.0, C. laxum v1.1, S.hybrid R570 v2.1(A), S.hybrid R570 v2.1(B), S.hybrid R570 v2.1(C), S.hybrid R570 v2.1(D), S.hybrid R570 v2.1(E), S.hybrid R570 v2.1(F), S.hybrid R570 v2.1(G), S.hybrid R570 v2.1(OS), M.sinensis v7.1 (A), M.sinensis v7.1 (B), Z.mays RefGen_V4 (A), Z.mays RefGen_V4 (B), Z.mays B73 v5 (A), Z.mays B73 v5 (B), P. vaginatum v3.1, U.fusca v1.1, S. viridis v4.1, S. italica v2.2, P. hallii v3.2.

When examining sorghum reference chromosomes, we found shared genes with other grass species such as SaccharumA (*Saccharum officinarum* x *spontaneum* var. R570), MisB (*Miscanthus sinensis), Setaria italica, Setaria viridis, Panicum virgatum, Urochloa fusca, Chasmanthium laxum, Oryza sativa* var. Nipponbare, and *Paspalum vaginatum* (Figure 5). This shared genetic heritage highlights the evolutionary connections within the Panicoideae group. Interestingly, in polyploid genomes like that of *Saccharum*, certain chromosomes maintain consistent genetic organization despite their complexity. For example, SaccharumD and SaccharumC exhibit stronger similarities to SaccharumA compared to other homologous chromosomes. Moreover, Miscanthus shares significant chromosomal similarities with *Sorghum* and *Saccharum*, emphasizing the interconnectedness of genetic structures across related species. Further exploration of polyploid genomes revealed intriguing dynamics, indicating past genetic recombination events and suggesting complex evolutionary paths for these organisms. Additionally, our examination of *Triticum aestivum* revealed a distinct genetic organization compared to other grass species, indicating unique evolutionary processes within this species. Similarly, *Zea mays* displays two distinct genetic patterns (Maize1 and Maize2), similar to *Miscanthus sinensis* (MisA and MisB), both of which have undergone significant recombination events. These findings underscore the complexity of grass genomes shaped by various evolutionary forces, including polyploidization and genomic rearrangements.

## Supporting information

Dias et al PGSGS Supplemental Table 1

Dias et al PGSGS Supplemental Table 2

Dias et al PGSGS Supplemental Table 3

## DATA AVAILABILITY

The supplementary tables associated with this study are available on Figshare and can be accessed via the following link: 10.6084/m9.figshare.26793190.v2. These data include a brief description of the contents, and the PGSGS source data. The pipeline used for this study, including additional scripts and documentation, is available as a supplemental resource on GitHub. The repository includes all relevant code for reproducing the analysis and details about the QuotaAlign pipeline, which was utilized for sequence alignment and comparative genomics. Users can access the repository on https://github.com/henrimdias/PGSGS

## AUTHOR CONTRIBUTIONS

HMD conceived and designed the bioinformatic approaches, analyzed the data, prepared figures and tables, authored or reviewed article drafts, and approved the final draft. GYAS analyzed the data, prepared figures and tables, reviewed the article, and approved the final draft. RVM and JVTR reviewed the supplementary analysis and article drafts and approved the final draft. JCS and MAVS conceived the project, discussed strategies and results, authored and reviewed drafts of the article, and approved the final draft.

## ACKNOWLEDGEMENTS

The authors extend their heartfelt gratitude to the dedicated teams at GaTE Lab and Schnable Lab for their unwavering motivation and profound curiosity in the realm of plant evolution. We also would like to express our deepest gratitude to the Holland Computing Center at the University of Nebraska-Lincoln (UNL) for their invaluable support and resources. The high-performance computing infrastructure and technical assistance provided by the center were crucial for the completion of this research. Their encouragement has been instrumental in propelling us forward in our pursuit of knowledge, recognizing the transformative impact that science and education have on lives. Special appreciation goes to Zach Shomo for providing unwavering support throughout the internship at the University of Nebraska-Lincoln. Their friendship has been instrumental in shaping a meaningful and enriching experience.

## TABLE AND FIGURES LEGENDS

**Supplementary Table S1. Assembly stats of 17 grass genomes publicly available on Phytozome v13**. This table provides comprehensive assembly statistics for grass genomes, sourced from Phytozome v13. The data includes details such as organism names, genome length, number of chromosomes, total scaffold length (in base pairs), number of scaffolds, minimum number of scaffolds containing half of the assembly (L50), shortest scaffold from the L50 set (N50), total contig length (in base pairs), number of contigs, minimum number of contigs containing half of the assembly (L50), shortest contig from the L50 set (N50), number of protein-coding transcripts, number of protein-coding genes, percentage of Eukaryote BUSCO genes, percentage of Embryophyte BUSCO genes, ploidy, and number of ploidy.

**Supplementary Table S2. Pan Grass Syntenic Gene Set (PGSGS) dataset**. This table presents a comprehensive collection of gene synteny and annotation data across multiple grass species. It includes information on whole syntenic and non-syntenic genes based on their relationship with Sorghum bicolor, along with corresponding PGSGS_IDs (Plant Genome Synteny Set IDs) and annotations from UniProt and Panther. Additionally, annotation enrichment analysis results are provided, obtained using eggNOG with a focus on the Poaceae reference. Annotations cover functional enrichments, biological insights, KEGG pathways, Gene Ontology (GO) terms, protein descriptions, Pfam domain, and other relevant annotations associated with genes within the Poaceae family.

**Supplementary Table S3. Comprehensive overview of enriched Gene Ontology (GO) terms within each syntelog cluster, encompassing both PGSGS-core syntelogs and variable syntelogs**. Detailed parameters from PANTHER and REVIGO analyses are included, with statistical corrections applied using False Discovery Rate (FDR) calculations. Graphical representations were constructed utilizing stringent criteria: REVIGO cutoffs for dispensability (≤ 0.05) and PANTHER cutoffs for fold-change (>1.5) alongside FDR thresholds (≤ 0.05). These refined parameters ensure robust identification and visualization of significant enrichments, enhancing the reliability and interpretability of the findings.

**Figure S1.**
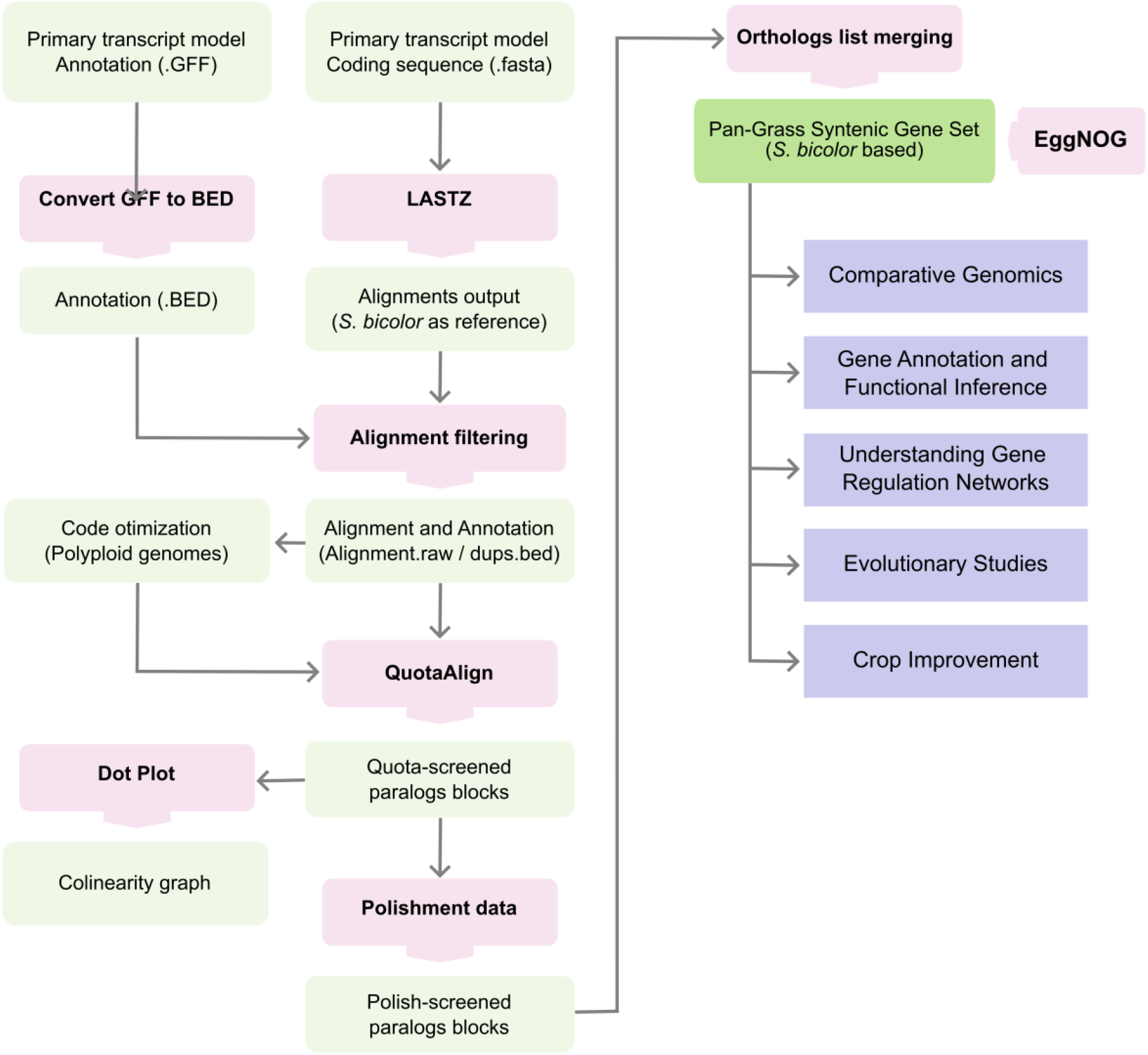
Workflow for comparative genomics and functional analysis of Pan-Grass Syntenic Gene Sets (PGSGS). The PGSGS analysis consists of seven major steps, some of which are further divided into sub-components. For each input file, a quality control is performed. This step includes both the application of filters and trimmers. (1) Primary transcript model annotation and coding sequence extraction. Input: .GFF and .fasta files containing the primary transcript model annotations and coding sequences. (2) Conversion and alignment. Convert GFF to BED: The GFF annotation file is converted to a BED format. LASTZ: Align coding sequences using *S. bicolor* as the reference genome, producing alignment outputs. (3) Alignment filtering and optimization. Filter alignments to refine results. Code Optimization: Improve the code for analyzing polyploid genomes. Generate alignment and annotation outputs. (4) QuotaAlign and paralog screening. QuotaAlign: Screen and filter for paralogous gene blocks. Polishment Data: Further refine paralogous blocks to enhance data quality. (5) Visualization and analysis. Dot Plot: Visualize syntenic blocks (collinearity graph). (6) Orthologs list merging and PGSGS building. Merge ortholog lists to form the PGSGS using *S. bicolor* as the base. (7) EggNOG enrichment. Integrate with the EggNOG database for additional functional annotations. (8) Downstream applications. Comparative Genomics: Compare genomic structures across species. Gene Annotation and Functional Inference: Annotate genes and predict their functions. Understanding Gene Regulation Networks: Analyze gene regulatory mechanisms. Evolutionary Studies: Investigate evolutionary relationships and patterns. Crop Improvement: Apply findings to enhance crop species.

